# Identification of a viral gene essential for the genome replication of a domesticated endogenous virus in ichneumonid parasitoid wasps

**DOI:** 10.1101/2024.01.18.576166

**Authors:** Ange Lorenzi, Fabrice Legeai, Véronique Jouan, Pierre-Alain Girard, Michael R. Strand, Marc Ravallec, Magali Eychenne, Anthony Bretaudeau, Stéphanie Robin, Jeanne Rochefort, Mathilde Villegas, Denis Tagu, Gaelen R. Burke, Rita Rebollo, Nicolas Nègre, Anne-Nathalie Volkoff

**Affiliations:** DGIMI, Montpellier University, INRAE, Montpellier, France; Department of Entomology, University of Georgia, Athens, Georgia, 30602, United States; INRAE, UMR Institut de Génétique, Environnement et Protection des Plantes (IGEPP), BioInformatics Platform for Agroecosystems Arthropods (BIPAA), Campus Beaulieu, 35042 Rennes, France; INRIA, IRISA, GenOuest Core Facility, Campus de Beaulieu, Rennes 35042, France; Univ Lyon, INRAE, INSA Lyon, BF2I, UMR 203, 69621 Villeurbanne, France

**Keywords:** Endogenous viral element, DNA amplification, *Hyposoter didymator*, Ichnovirus, polydnavirus, viral replication, RNA interference, co-option, co-evolution

## Abstract

Thousands of endoparasitoid wasp species in the families Braconidae and Ichneumonidae harbor “domesticated endogenous viruses” (DEVs) in their genomes. This study focuses on ichneumonid DEVs, named ichnoviruses (IVs), which derive from an unknown virus and produce virions in ovary calyx cells during the pupal and adult stages of female wasps. Females inject IV virions into host insects when laying eggs. Virions infect cells which express IV genes with functions required for wasp progeny development. IVs have a dispersed genome consisting of two genetic components: proviral segment loci that serve as templates for circular dsDNAs that are packaged into capsids, and genes from an ancestral virus controlling virion production. Because of the lack of homology with known viral genes, the molecular control mechanisms of IV genome are largely uncharacterized. We generated a chromosome-scale genome assembly *for Hyposoter didymator* and identified a total of 67 *H. didymator* ichnovirus (HdIV) loci distributed across the 12 wasp chromosomes. By analyzing genomic DNA levels, we found that all HdIV loci were locally amplified in calyx cells during the wasp pupal stage, suggesting the implication of viral proteins in DNA replication. We tested a candidate HdIV gene, *U16*, encoding a protein with a conserved domain found in primases and which is transcribed in calyx cells during the initial stages of replication. Knockdown of *U16* by RNA interference inhibited amplification of all HdIV loci, as well as HdIV gene transcription, circular molecule production and virion morphogenesis in calyx cells. Altogether, our results showed that viral DNA amplification is an early step of IV replication essential for virions production, and demonstrated the implication of the viral gene *U16* in this process.

**Author Summary:** Parasitoid “domesticated endogenous viruses” (DEVs) provide a fascinating example of eukaryotes acquiring new functions through integration of a virus genome. DEVs consist of multiple loci in the genomes of wasps. Upon activation, these elements collectively orchestrate the production of virions or virus-like particles that are crucial for successful parasitism of host insects. Despite the significance of DEVs for parasitoid biology, the mechanisms regulating key steps in virion morphogenesis are largely unknown. In this study, we focused on the ichneumonid parasitoid *Hyposoter didymator*, which harbors an ichnovirus consisting of 67 proviral loci. Our findings reveal that all proviral loci are simultaneously amplified in ovary calyx cells of female wasps during the early pupal stage suggesting a hijacking of cellular replication complexes by viral proteins. We tested the implication of such a candidate, *U16*, encoding a protein with a weakly conserved primase C-terminal domain. Silencing *U16* resulted in inhibited viral DNA amplification and virion production, underscoring the key role of this gene for ichnovirus replication. This study provides evidence that genes involved in viral DNA replication have been conserved during the domestication of viruses in the genomes of ichneumonid wasps.

## Introduction

Endogenous viral elements (EVEs) refer to viral sequences in eukaryotic genomes that originate from complete or partial integration of a viral genome into the germline [1]. While retroviruses are the best-known sources of EVEs, bioinformatic studies have also identified non-retroviral EVEs across a diverse range of organisms [2]. Although many EVEs become non-functional and decay through neutral evolution [3], some have been preserved and repurposed by their hosts for new functions, often as short regulatory sequences or individual genes [4,5]. A notable exception to this pattern is observed in domesticated endogenous viruses (DEVs) that have been identified in four lineages of endoparasitoid wasps - insects that lay eggs and develop within the bodies of other insects [6]. Parasitoid DEVs consist of numerous genes conserved within the wasp genome that originate from the integration of complete viral genomes. Unlike other EVEs, these genes remain functional and actively interact to produce virus particles in calyx cells, which are located in the apical part of the oviducts of female wasps [7]. Viral particles are produced in the pupal and adult stages, and accumulate in the oviducts of the wasp. Adult female wasps inject these particles along with eggs into insect hosts where they have essential functions in the successful development of wasp offspring [8].

Parasitoid DEVs are prevalent among species in two wasp families named the Braconidae and Ichneumonidae. The DEVs identified in these families have evolved from different virus ancestors but through convergence have been similarly repurposed to produce either virions containing circular double-stranded (ds) DNAs or virus-like particles (VLPs) lacking nucleic acid. The hyperdiverse Microgastroid complex in the family Braconidae harbors DEVs named bracoviruses (BVs). BVs evolved from a virus ancestor in the family Nudiviridae [9]. Wasps harboring BVs produce virions containing circular dsDNAs. Other braconids in the subfamily Opiinae and ichneumonids in the subfamily Campopleginae independently acquired two other distinct nudiviruses that wasps have coopted to produce VLPs [10, 11]. The fourth identified DEV lineage, named ichnoviruses (IVs), is present in two ichneumonid subfamilies (Campopleginae and Banchinae) which produce virions containing circular dsDNAs. Unlike the other three DEVs, IVs likely originated from a Nucleocytoplasmic Large DNA Virus (NCLDV) but the precise ancestor remains unknown [12, 13].

BVs have been more studied than IVs but the latter are intriguing because of their uncertain origins. Despite differences in ancestry and gene content, BV and IV genomes are similarly organized into two components that have distinct functions [14]. Insights into the genome components of IVs primarily derive from sequencing two campoplegine wasps named *Hyposoter didymator* and *Campoletis sonorensis* [15], along with calyx transcriptome studies [12, 13, 16, 17] and proteomic analyses of purified virions [12, 13]. The first genome component of IVs are domains in the wasp genome that show evidence of deriving from the virus ancestor and having essential functions in virion formation. These domains, named “Ichnovirus Structural Protein Encoding Regions” (IVSPERs), contain intronless genes that are specifically transcribed in calyx cells [12, 13, 17]. Most IVSPER genes are transcribed at the onset of pupation in hyaline stage 1 pupae [16], and some genes in IVSPERs encode proteins associated with IV virions [12, 13]. Six genes have been knocked down by RNA interference (RNAi) in *H. didymator* which demonstrated that they have functions in virion assembly or cell trafficking [16]. Five IVSPERs have been identified in the *H. didymator* and *C. sonorensis* genomes [15], while three have been identified in the genome of the more distantly related banchine *G. fumiferanae* [13]. The content of IVSPER genes is notably similar between ichneumonid wasp species [12, 13, 17], and their gene order is well-conserved among campoplegine species [15]. Additionally, one intronless gene (*U37*) was identified in the *H. didymator* and *C. sonorensis* genomes outside of any IVSPER with features suggesting it also derives from the virus ancestor [15]. Together, these genes, whether found within or outside IVSPERs, represent the fingerprints of the ancestral viral machinery essential for virion production and are designated as IV core replication genes. Notably, none of these genes are packaged in virions, indicating that IV core genes can only be transmitted vertically through the germline of associated parasitoids.

The second component of IV genomes are domains referred to as “proviral segments,” which are amplified in calyx cells and produce the circular dsDNAs that are packaged into capsids [18, 19]. The number of proviral segments, typically exceeding 50, are widely dispersed in wasp genomes and exhibit considerable variability between wasp species, [15]. Each proviral segment is characterized by flanking direct repeats (DRs) of variable length (<100 bp to >1 kb) and homology that identify where homologous recombination processes occur to produce circularized DNAs [18, 19]. Some IV proviral segments also contain internal repeats that facilitate additional homologous recombination events, and produce multiple overlapping or nested circularized DNAs per proviral segment [15, 18]. Proviral segments encode genes with and without introns that are predominantly expressed in the hosts of wasps after virion infection [20, 21, 22, 23]. While IV core replication genes represent the conserved viral machinery that produces virions in calyx cells, proviral segments constitute the IV genome components that virions transfer to the hosts wasps parasitize. These segments also play a major role in the virulence of IVs, which contributes to the successful development of parasitoid progeny.

The replication of IVs, encompassing the processes leading to the production of virions containing IV segments, occurs within the nuclei of calyx cells during pupal and adult developmental stages [7, 24]. Electron microscopy studies of *H. didymator* ichonovirus (HdIV) shows that fusiform-shaped capsids are individually enveloped in the nuclei of calyx cells during the late pupal stage (pigmented pupae, stage 3) [16]. These enveloped “subvirions” exit the nucleus, traverse the cytoplasm, and exit calyx cells by budding, resulting in mature virions with two envelopes that accumulate in the calyx lumen of the ovaries [7, 24]. Earlier findings indicated that IVSPERs and proviral segments undergo amplification in newly emerged adult wasps [12]. However, these data focused on only a subset of IVSPER genes and one proviral segment, leaving our knowledge of whether all IV genome components are amplified in calyx cells incomplete. Similarly, the initiation time of amplification during pupal development and IV virion production remains unknown. The specific role of IV core genes in virion production is also poorly documented when compared to BVs [25, 26]. The limited sequence homology of IVSPER genes with genes in other viruses provides minimal insights into potential functions. To date, only the six genes mentioned above that are involved in subvirion assembly or cell trafficking have been studied [16].

In this work, we explored IV replication using the campoplegine wasp *H. didymator*. We first generated a chromosome-level assembly for the *H. didymator* genome. Through this assembly, we determined that all genome components undergo local amplification in calyx cells which initiates between pupal stages 1 and 2. Notably, IVSPERs, isolated IV core genes, and proviral segments were amplified in large regions with non-discrete boundaries. Next, we studied the function of *U16* which is located on *H. didymator* IVSPER-3. *U16* is one of the most transcribed IVSPER genes during the initial pupal stage and contains a weakly conserved domain found in the C-terminus of primases. RNAi knockdown of *U16* inhibited virion formation. Knockdown also significantly reduced DNA amplification of all HdIV genome components, which decreased transcript abundance of IV core genes and the abundance of circular dsDNA viral molecules. We conclude *U16* is an essential gene for amplification of the HdIV genome and virion production, demonstrating that genes from the IV ancestor regulating IV replication have been conserved during virus domestication. Additionally, our results show that viral DNA amplification is essential for IV virion production.

## Results

### Genomic localization of *Hyposoter didymator* IV components in a novel chromosome-level assembly

The genome assembly for H. didymator we previously generated [15] consisted of 2,591 scaffolds with an N50 of 4 Mbp. We concluded this assembly was overly fragmented to evaluate DNA amplification in calyx cells during virion morphogenesis. We therefore used proximity ligation technology to produce a new chromosome level assembly consisting of twelve large scaffolds that corresponds with the haploid karyotype for H. didymator [27]. The sizes of these scaffolds ranged from 6.7 Mbp to 29.3 Mbp (S1 Dataset A, B).

The five IVSPERs (IVSPER-1 to IVSPER-5), the predicted IV core gene (*U37*) located outside of an IVSPER, and 53 of the 54 previously identified proviral segment loci (Hd1 to Hd54) [15] were identified in the new assembly. The new assembly did not include the scaffold containing Hd51, possibly due to low-quality sequencing data (S1 Dataset, B). Our chromosome-level assembly revealed that each scaffold contained at least one HdIV locus, but notably, all IVSPERs and 40% of the proviral segment loci resided on two (scaffold 7 and 11) (S1 Dataset, B).

While three IVSPERs and the majority of proviral segments were distantly located from each other in the *H. didymator* genome, there were exceptions to this pattern including certain pairs of proviral segments separated by less than 20 kb (e.g., Hd36 and Hd38; Hd46 and Hd43; Hd44.1 and Hd44.2; Hd12 and Hd16). In all of these cases, the paired segments exhibited significant homology which suggested they derive from recent duplication events (S1 Dataset, C). Additionally, several proviral segments were in proximity to IVSPERs or IV replication genes that resided outside of IVSPERs (e.g., Hd46 near U37; Hd29 and Hd24 on each side of IVSPER-2; Hd15 near IVSPER-1; also see below).

### Amplification of *Hyposoter didymator* IV genome components in calyx cells during wasp pupal development

To investigate whether all or only specific components of the HdIV genome undergo amplification in association with virion morphogenesis, we isolated DNA from calyx cells from stage 1 pupae (one day old, hyaline) and stage 3 pupae (five days old, pigmented abdomen). We then generated paired-end libraries, which were sequenced using the Illumina platform, followed by read alignment to the new chromosome-level genome assembly. When analyzing the reads from stage 1 pupae, read coverage per HdIV locus did not differ significantly from the coverage of randomly selected regions of the same size from the rest of the wasp genome (Fig 1A). In contrast, read coverage for stage 3 pupae was higher for all HdIV loci when compared to the rest of the wasp genome or to values obtained for pupal stage 1 (Fig 1A, S1 Table).

**Fig 1.**
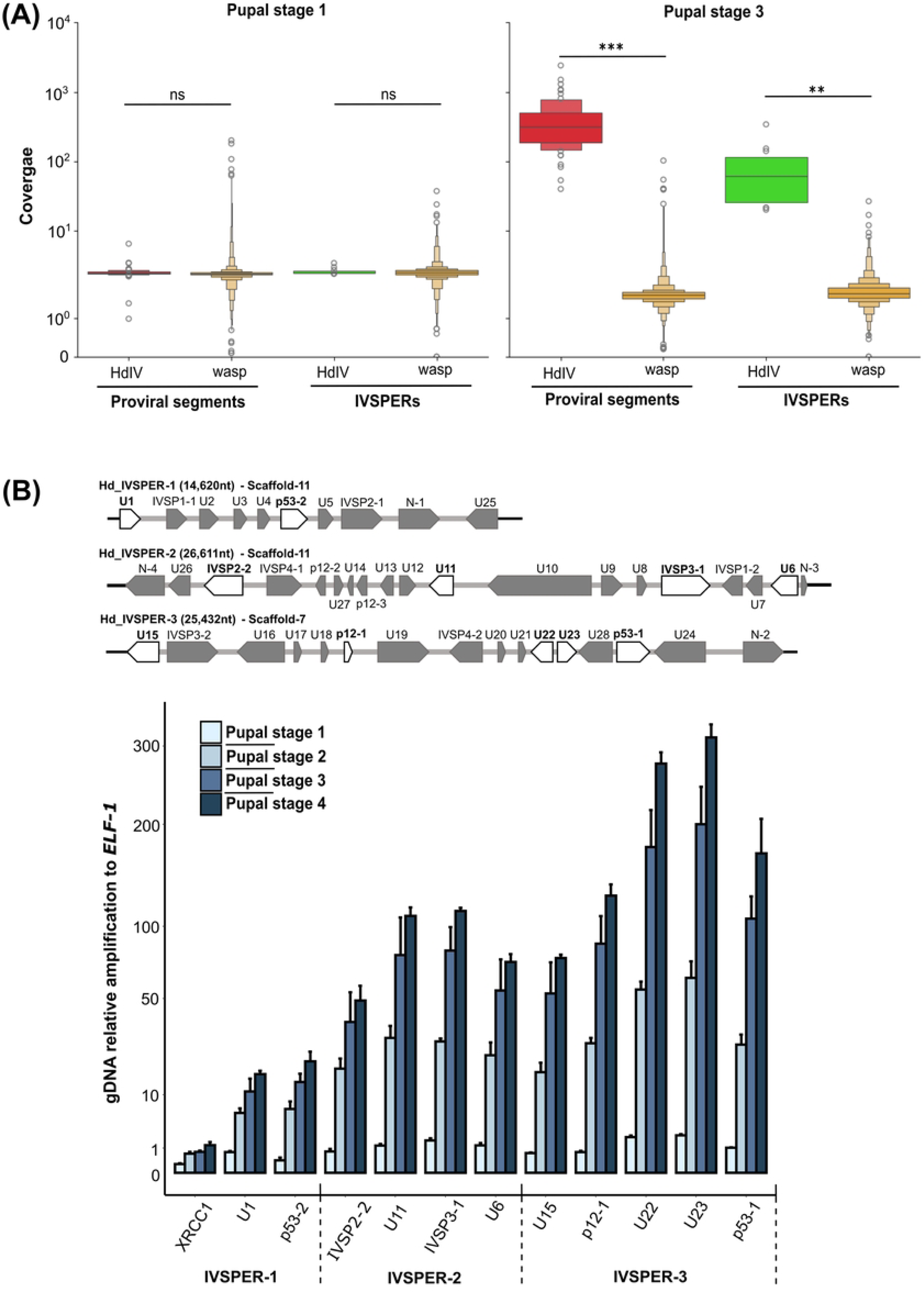
DNA amplification of HdIV loci. **(A)** Coverage of HdIV loci compared to the rest of the wasp genome. Read coverage values per analyzed region (see Materials and Methods) are presented for each locus type (proviral segments and IVSPERs) at pupal stage 1 (hyaline pupa) and pupal stage 3 (pigmented pupa). The coverages per HdIV locus are compared to the coverage per random genome regions outside of HdIV loci (wasp). Note that the coverage value for random wasp regions is lower for DNA samples collected from stage 3 versus stage 1 pupae. This difference is attributed to the higher proportion of reads mapping to HdIV regions among the total number of reads in stage 3 compared to stage 1. The significance levels are indicated as follows: ns = non-significant, **p<0.01, and ***p<0.001. **(B**) qPCR analysis of select IVSPER genes in calyx cells during wasp pupal development. Top panel. A schematic representation of *H. didymator* IVSPERs-1, -2, and -3 (GenBank GQ923581.1, GQ923582.1, and GQ923583.1); genes selected for qPCR assays are highlighted in white. U1-24 are unknown protein-encoding genes, while IVSPs are members of a gene family encoding ichnovirus structural proteins. Bottom panel. Genomic (g) DNA amplification levels of IVSPER genes and wasp *XRCC1* in calyx cells from pupal stage 1-4. The XRCC1 (X-Ray Repair Cross Complementing 1) encoding gene is located 1,200 bp from *U1* (position 3,270,470 to 3,272,519 in Scaffold-11). Data corresponds to gDNA amplification relative to amplification of the housekeeping gene elongation factor 1 (ELF1). The Y-axis was transformed using the square root function for better data visualization.

To more precisely investigate the temporal dynamics of amplification, we conducted relative quantitative (q) PCR assays that measured copy number of genes in IVSPER-1, -2, and -3 in calyx DNA samples that were collected from stage 1-4 pupae. We compared these treatments to DNA samples from hind legs of stage 1 pupae where no HdIV replication occurs. We also included a wasp gene (*XRCC1*) located in close proximity to IVSPER-1. Results showed that copy number of each tested gene was similar in calyx and hind legs in stage 1 pupae, indicating none were amplified during the initial pupal stage. Subsequently, the copy number of each gene increased progressively with each pupal stage (Fig 1B). While exhibiting lower amplification levels than the IVSPER genes we analyzed, a similar trend was observed for the wasp gene *XRCC1* (Fig. 1B). These findings indicated IVSPER amplification in calyx cells begins between pupal stage 1 and stage 2, which further increased in pupal stage 3 and 4.

### Differential levels of amplification across all components of the HdIV genome

The qPCR results presented in Fig 1 indicated amplification levels varied, with genes in IVSPER-3 exhibiting higher levels of amplification than genes in IVSPER-1 and -2 (Fig 1B). This variability was corroborated genome-wide by analyzing read coverage per position and the ratio between stage 3 and stage 1 (Fig 2, S1 Fig). Amplification levels of IVSPER loci, determined at the summit of the coverage curve, ranged from 10X for IVSPER-5 in Scaffold-7 to over 200X for IVSPER-3 in Scaffold-3 (S1 Table). This observation aligned with the findings from qPCR analyses, indicating that genes in IVSPER-3 were more highly amplified than those in IVSPER-1 and -2 (Fig 1B). Read mapping further indicated that the peak of amplification occurs toward the middle of each IVSPER (Fig 1B, S1 Fig), consistent with qPCR analyses revealing that within each IVSPER, genes closer to the cluster boundary tended to exhibit lower levels of amplification compared to genes situated in the middle of the cluster (Fig 1B).

**Fig 2.**
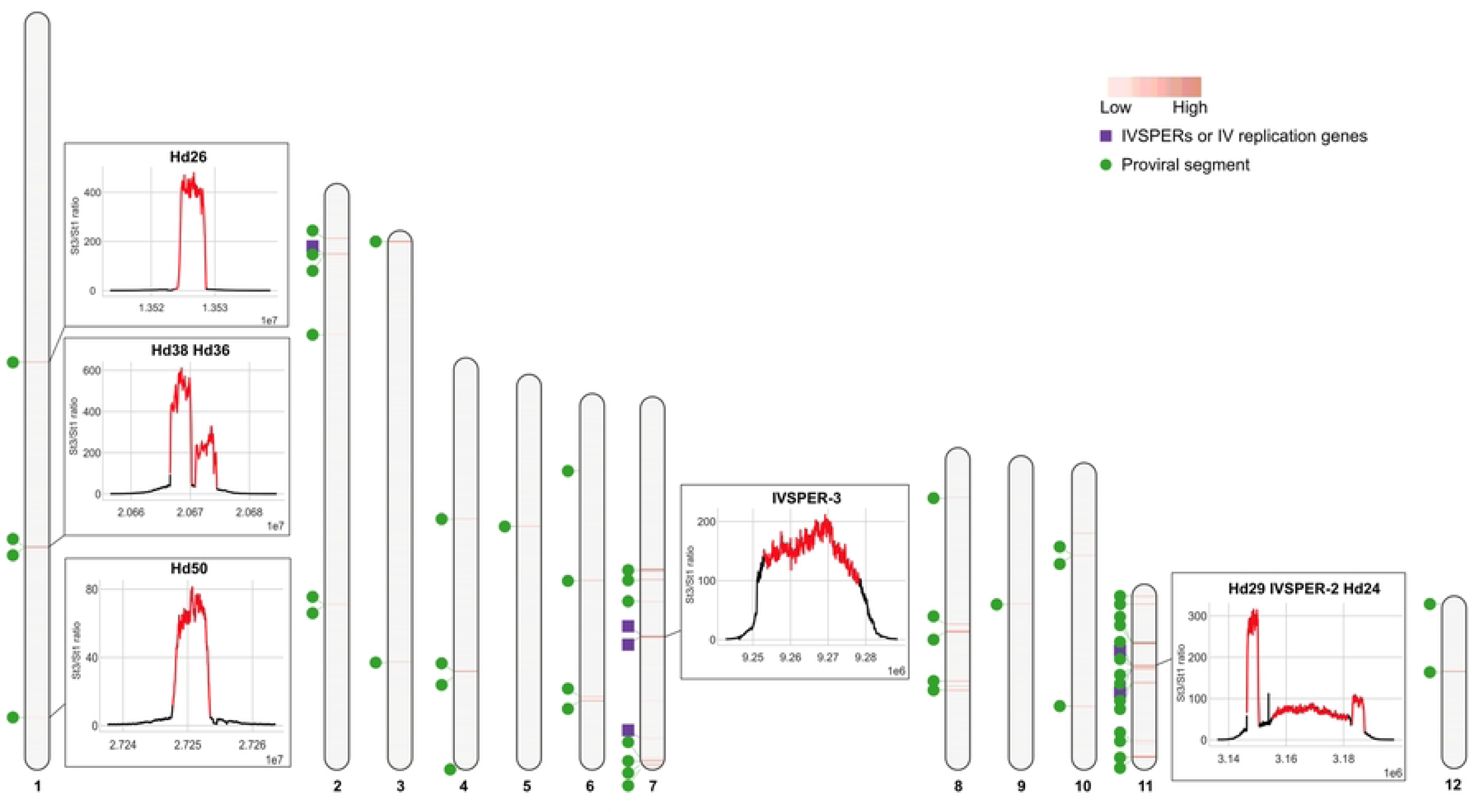
HdIV DNA amplification. DNA amplification in pupal stage 3 was assessed by mapping genomic DNA Illumina reads against the 12 large *H. didymator* genome scaffolds. In each scaffold, red bars indicate amplified loci, with the intensity of red corresponding to increased values of the CPM ratio between pupal stage 3 and pupal stage 1. The positions of IVSPERs and isolated IV replication genes are indicated by purple squares, while proviral segments are indicated by green circles. For selected HdIV loci, amplification curves (representing the ratio of the CPM values calculated for 10 bp intervals between pupal stage 3 and pupal stage 1) are shown in boxes. Amplification curves for all of the annotated HdIV loci are shown in S1 Fig. Each HdIV locus is indicated in red while 10,000 bp of flanking sequence on each side of the locus is also shown. For proviral segments, loci are defined as the sequence delimited by two direct repeats; IVSPERs are defined as the region between the start and stop codon of the first and last coding sequences in the cluster; isolated IV replication genes are defined by their coding sequence.

Proviral segment loci were relatively more amplified than IV replication gene loci, and also variable in intensity (Fig 2, S1 Fig). For example, coverage ratio between stages 3 and 1 ranged from 30X for proviral locus Hd40 in Scaffold-6 to over 1,100X for Hd27 in Scaffold-7 (S1 Table) at the summit of the coverage curves. Variability in the number of reads mapping to a given proviral locus was consistent with earlier studies indicating that the circularized DNAs packaged into IV capsids are non-equimolar in abundance [8, 28].

All proviral segments consistently exhibited a substantial increase in amplification that peaked between the two DRs (as exemplified by Hd14 or Hd12 in S2 Fig). For numerous proviral loci, the reads mapping between the flanking DRs displayed uniform coverage. However, in other cases, peaks with varying read coverage were evident (as exemplified by Hd32 or Hd16 in S2 Fig). This differential coverage usually applied to proviral segments containing more than one pair of DRs, as illustrated by proviral locus Hd11 (Fig 3A) or Hd32 and Hd16 (S2 Fig). Previous studies indicated Hd11 contains two pairs of DRs, enabling the formation of two nested, circularized segments termed Hd11-1 (formed by recombination between DR1Left (DR1L) and DR1Right (DR1R)) and Hd11-2 (formed by recombination between DR2L and DR2R) (Fig 3A). Reads mapping to the Hd11 locus (bounded by DR1L and DR2R) exhibited three relatively uniform plateaus of different values. Two plateaus corresponded to reads mapping to the predicted locations of Hd11-1 (235X) and Hd11-2 (111X), while the central region with higher coverage (311X) corresponded to reads mapping to both nested segments (Fig 3A). This differential coverage would not be expected if reads mapped only to Hd11 chromosomal DNA. Consequently, the pattern of proviral segment coverage suggested part of the coverage values were due to reads mapping to amplification intermediates and/or circularized dsDNAs that were also present in our DNA samples. Some amplified HdIV loci contain both an IVSPER and proviral segments. Two of these loci resided on Scaffold-11 (Hd29, IVSPER-2, Hd24, and Hd33, Hd15, IVSPER-1 (Fig 3B)). For these loci, the amplification curves spanned the length of the amplified region (yellow dotted line in Fig 3B) but were interrupted by peaks corresponding to the length of proviral segments. This pattern suggested amplification levels of the chromosomal form of the proviral segments could correspond to the IVSPER amplification curves, but were higher because reads additionally mapped to circular dsDNAs or amplification intermediates.

**Fig 3.**
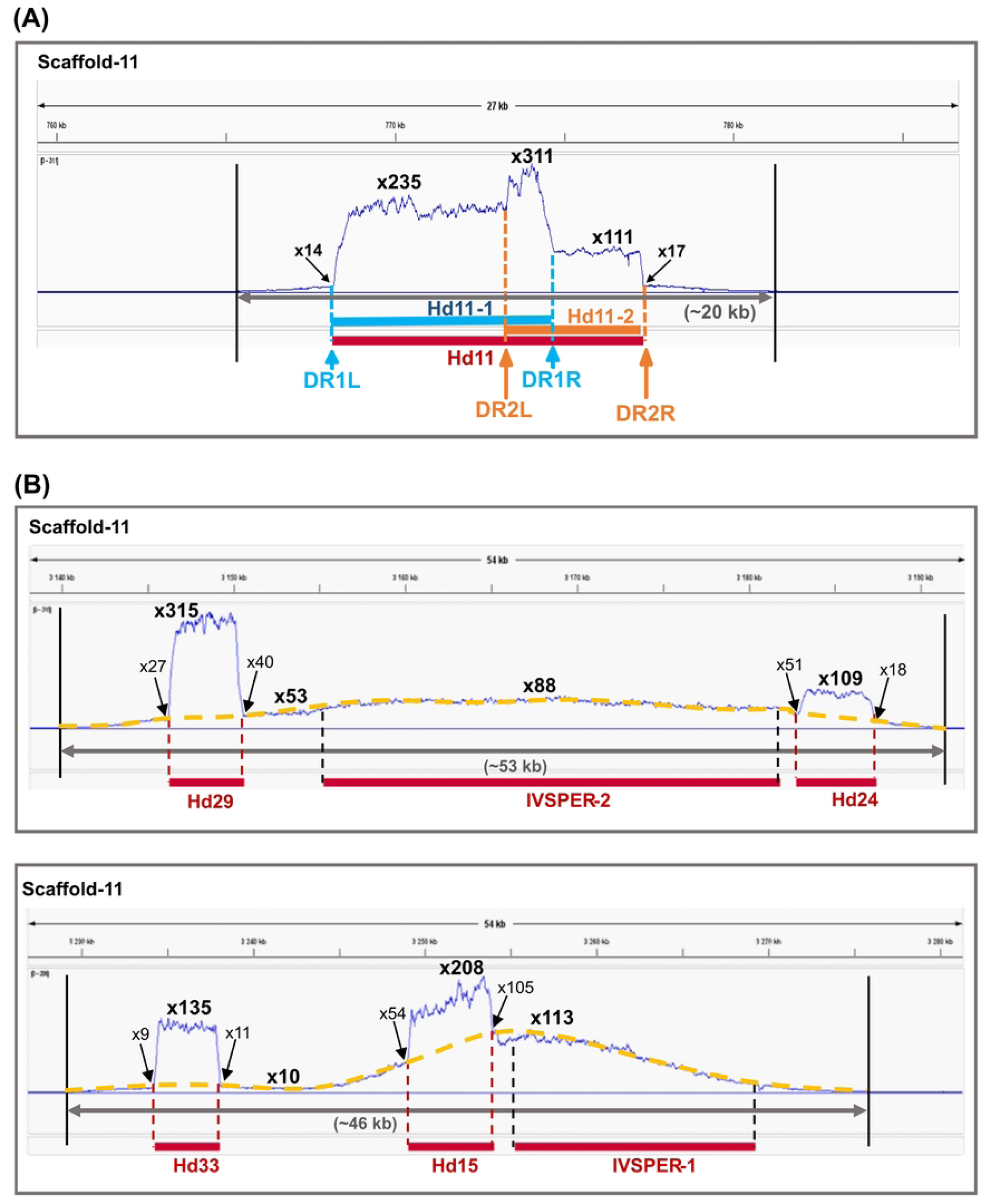
HdIV amplified regions in Scaffold-11. **(A)** Detail of the amplified region at the Hd11 locus. **(B)** Detail of two other amplified regions containing IVSPERs and HdIV proviral loci. In **(A)** and **(B)**, amplification curves represent the ratio of the CPM values (calculated for 10 bp intervals) obtained in pupal stage 3 compared to pupal stage 1. For each locus, amplification values at the summit of the peaks and at the start and end positions of HdIV segments are indicated. In **(B)**, amplification curves of IVSPERs are highlighted in yellow. Each amplification curve figure was generated by Integrated Genome Viewer (IGV) [29].

### Amplification of *H. didymator* IV genome components in extensive wasp genome domains with undefined boundaries

Since our read coverage data indicated amplified regions were larger than the annotated HdIV loci (Fig 2, S1 Fig), we used the MACS2 peak calling program, originally developed for chromatin immunoprecipitation sequencing experiments, to identify areas in the *H. didymator* genome that were enriched for reads when compared to a control [30]. Amplification peaks were called with MACS2 using alignments from stage 3 pupae as the treatment and alignments from stage 1 pupae as the control. MACS2 identified all HdIV genome components that we had annotated in our earlier study [15] plus several previously unrecognized domains (S2 Table). Manual curation (see Materials and Methods section) indicated three of these new domains were proviral segment loci that we named Hd52, Hd53, and Hd54. Five others were intronless genes, suggesting origins from the IV ancestor, that were outside of IVSPERs. We thus named these genes *U38, U39, U40, U41*, and *U42*. The remaining domains detected by MACS2 either contained predicted wasp genes or lacked any features that identified them as IV replication genes or proviral segments. Altogether, the MACS2 algorithm predicted a total of 55 domains in the *H. didymator* genome containing HdIV loci. Two proviral segments (Hd45.1 on Scaffold-4 and Hd2-like on Scaffold-7) escaped MACS2 detection, possibly because they were located too close to the ends of each scaffold. However, our read mapping data clearly indicated these two segments are amplified in stage 3 (Table 1) with a profile similar to the other segments (S1 Fig). In total, our read mapping and MACS2 data indicated the *H. didymator* genome contains 67 HdIV loci (56 proviral segments, five IVSPERs, and six predicted IV replication genes that reside outside of IVSPERs) that are amplified in calyx cells at pupal stage 3 (Fig 2, Table 1).

**Table 1.** All HdIV loci amplified in calyx cells from stage 3 pupae identified by read mapping and/or the MACS2 algorithm. For each scaffold, the position and size of the HdIV loci are indicated. Loci newly identified in the present work are marked with asterisks. Corresponding amplified regions (i.e., the peak predicted by the MACS2 algorithm) are provided for each locus or groups of loci. Start and end positions delimiting the HdIV loci and the amplified regions detected by MACS2 are indicated. The distance between the start or the end of the amplified region and the locus is presented. For each HdIV locus and amplified region detected by MACS2, coverage values are provided for calyx cell samples collected from stage 1 or stage 3 pupae. Coverage is based on the length of the HdIV locus or amplified region. ND indicates amplified regions not detected by MACS2.

Our overall results also indicated all amplified regions in the *H. didymator* genome containing HdIV loci consist of the annotated HdIV locus plus flanking wasp sequence consistent with our detailed analysis of the wasp gene *XRCC1* that is located in close proximity to IVSPER-1 (Fig 1B). Across all HdIV loci, we determined that the flanking regions containing wasp sequence that were amplified varied from 7,000 to 15,000 bp (Table 1). The total size of the amplified regions ranged from 10,692 bp (Hd28 on Scaffold-12) to 54,005 bp (IVSPER-2 on Scaffold-11). Most amplified regions contained a single HdIV locus, but seven contained a mix of HdIV genome components (Table 1). Three amplified regions contained the neighboring and closely related proviral segments mentioned above (e.g., Hd36 and Hd38 on Scaffold-1, Hd44.1 and Hd44.2 on Scaffold-2, Hd12 and Hd16 on Scaffold-11). In addition to the two examples noted above on Scaffold 11 (see Fig. 3B), two other amplified loci also contained both IVSPERs and proviral segments (U37, Hd46, and Hd43 on Scaffold-2; U40 and Hd39 on Scaffold-9). Lastly, we searched for sequence signatures that potentially identify the amplification boundaries for each HdIV locus. However, our analysis identified only low-complexity A-tract sequences, which were not specific to HdIV components as they were also found in random wasp genomic sequences (S3 Fig). Thus, no motifs were identified that distinguished the amplification boundaries of HdIV loci.

### RNAi knockdown of *U16* inhibits virion morphogenesis

We selected the gene *U16* located on *H. didymator* IVSPER-3 as a factor with potential functions in activating IV replication. *U16* is conserved among all IV-producing wasps for which genome or transcriptome data is available (Fig 4A). In *H. didymator* calyx cells, *U16* is also one of the most transcribed IV genes detected in calyx cells from stage 1 pupae [16]. Sequence analysis using the basic local alignment search tool and DeepLoc2.0 predicted all U16 family members contain a C-terminal alpha-helical domain (PriCT-2) of unknown function that is present in several primases [31] (Iyer et al., 2005) and a nuclear localization signal (Fig 4A, S2 Dataset). We next assessed the effects of knocking down *U16* by RNAi on virion morphogenesis in calyx cells. We injected newly pupated wasps with dsRNAs that specifically targeted *U16* using previously established methods [16]. RT-qPCR analysis indicated transcript abundance in the calyx of newly emerged adult females was reduced more than 90% when compared to control wasps that were injected with ds*GFP* (Fig 4B). Inspection of the ovaries further indicated that the calyx lumen of control wasps contained blue ’calyx fluid’ indicative of HdIV virions being present, whereas almost no calyx fluid was seen in ds*U16*-injected wasps (Fig 4B). Examination of calyx cell nuclei by transmission electron microscopy similarly showed that calyx cells in one day old control females contained an abundance of subvirions, whereas no subvirions were observed in treatment wasps (Fig 4C). We thus concluded that U16 is required for virion morphogenesis.

**Fig 4.**
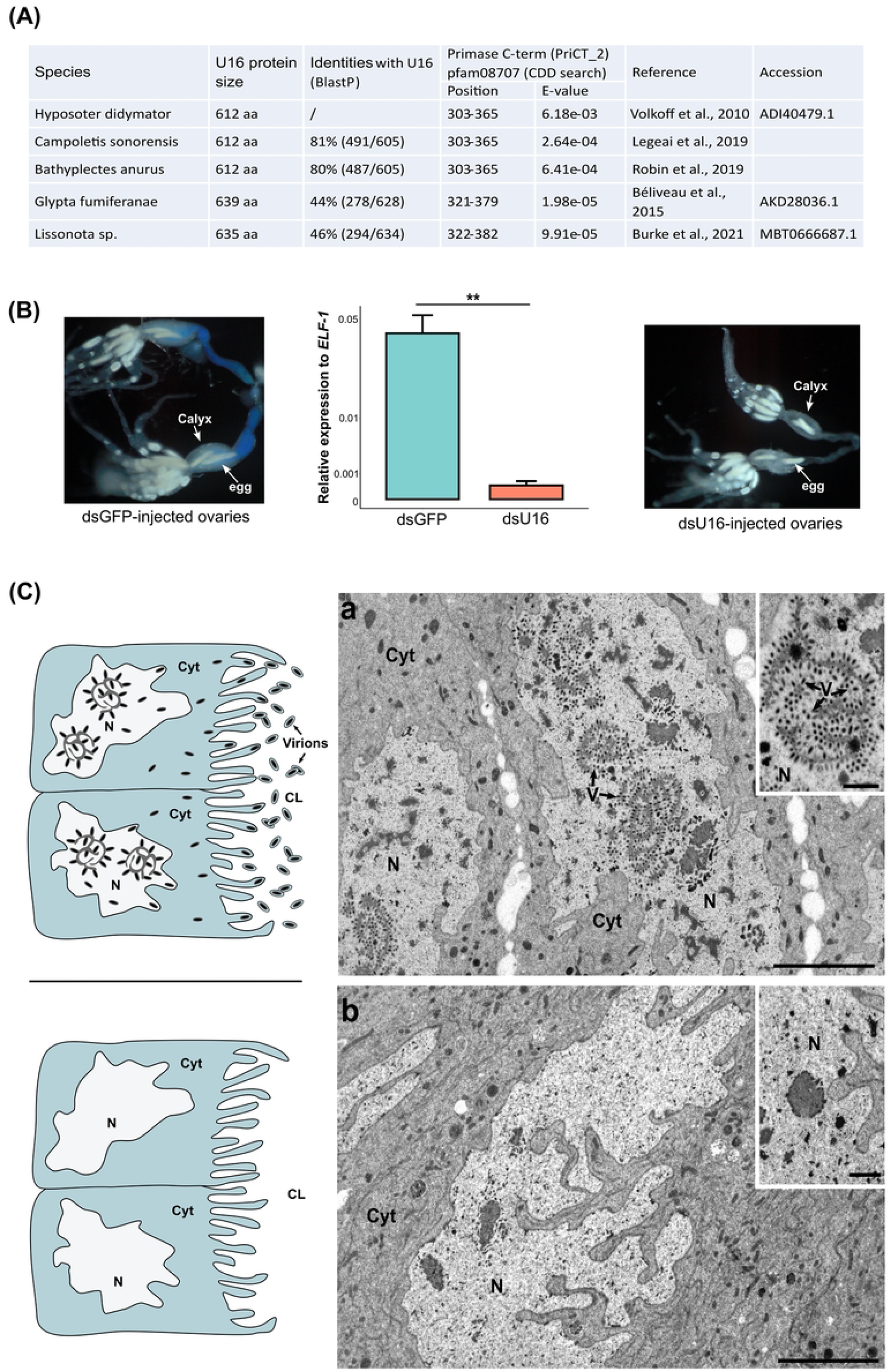
RNAi knockdown of *U16.* **(A)** U16 proteins identified in the campoplegine *Hyposoter didymator* [12], *Campoletis sonorensis* [15], and *Bathyplectes anurus* [17], and in two banchine wasps *Glypta fumiferanae* [13] and *Lissonota* sp. [32]. For each, protein size, percentage of identity with *H. didymator* protein and location of the PRiCT_2 domain are indicated. **(B)** RT-qPCR data showing relative expression of *U16* in ds*GFP* (control) and ds*U16* injected females. ** p<0.01. Images of ovaries dissected from newly emerged adult females that were injected with ds*GFP* (left) or ds*U16* (right). Note the blue color in the oviduct of the ds*GFP* control indicating the presence of HdIV virions. **(C)** Schematics and electron micrographs showing that (a) calyx cell nuclei (N) from females treated with ds*GFP*-injected contain subvirions (V) while (b) calyx cell from a ds*U16*-injected wasps do not. This results in no accumulation of virions in the calyx lumen as illustrated in the schematic images. CL, calyx lumen; Cyt, cytoplasm. Scale bars = 5 μm, zooms = 1 μm.

### RNAi knockdown of *U16* also disables amplification of HdIV loci

Since *U16* contained a domain found in primases, we investigated whether RNAi knockdown also disabled amplification of HdIV genome components. We injected newly pupated wasps with ds*U16* or ds*GFP*, followed by isolation and deep sequencing of calyx cell DNA from stage 3 pupae in three independent replicates. Mapping the reads from ds*GFP*-treated calyx samples to the *H. didymator* genome indicated all HdIV loci were amplified as evidenced by higher coverage values when compared to random regions of the wasp genome (Fig 5A). Conversely, coverage values did not differ between HdIV loci and other regions of the wasp genome in ds*U16-*treated calyx samples (Fig 5A). When analyzing coverage per each HdIV genome component (IVSPERs, isolated IV replication genes, or HdIV proviral segments), we also determined that values were systematically lower for the ds*U16* than ds*GFP*-treatments (Fig 5B and 5C, S3 Table).

**Fig 5.**
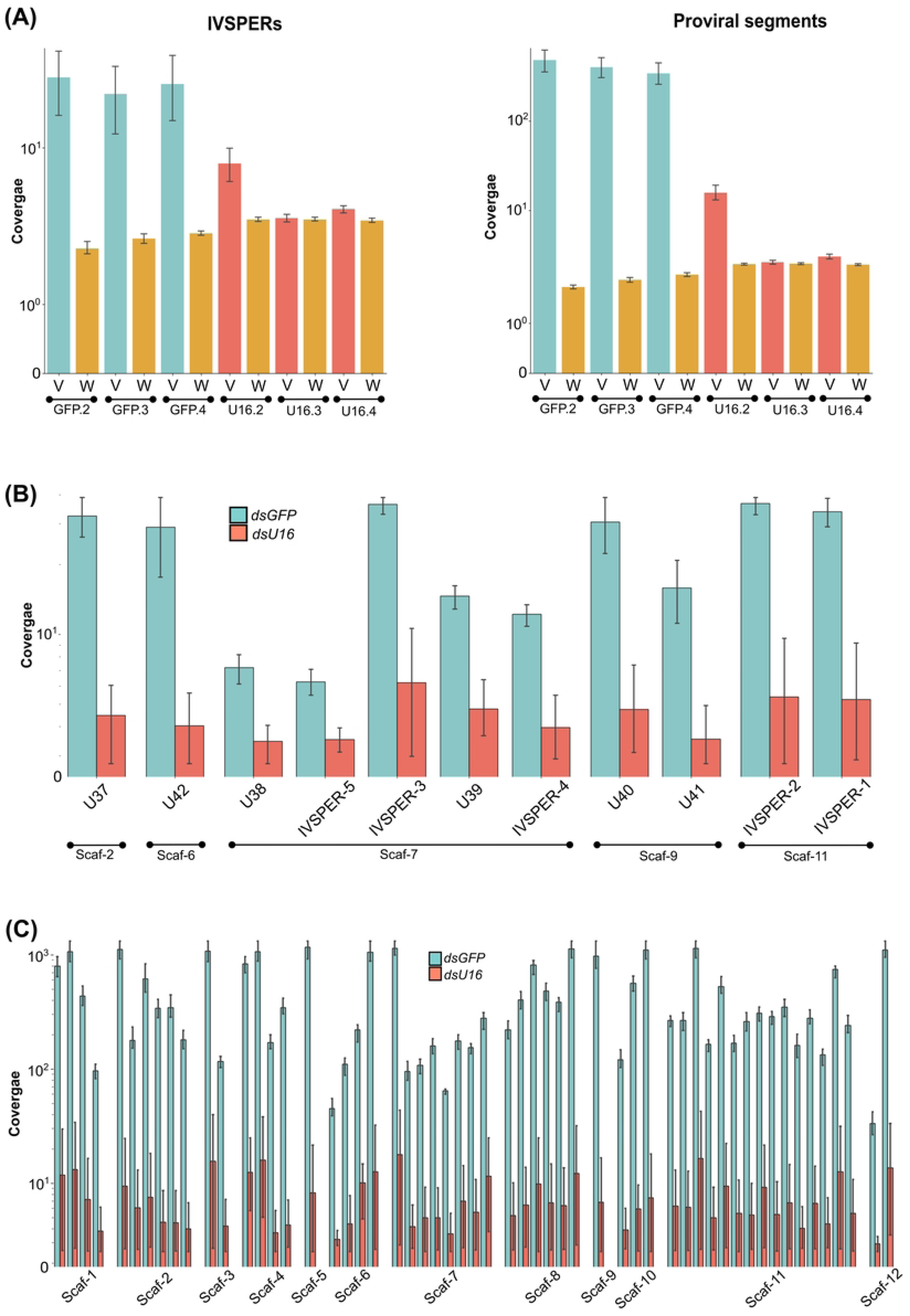
Impact of *U16* RNAi knockdown on DNA proviral amplification. **(A)** Comparative distribution of read coverages in ds*GFP*- and ds*U16*-injected females. For each of the three replicates, coverage values are given per HdIV loci (V) and per random genome regions outside of the HdIV loci (W), both with the same size distribution. IVSPERs and IV replication genes loci are shown in the left panel, while proviral segment loci are shown in the right panel. **(B)** Coverage values per IVSPERs, and per IV replication genes residing outside an IVSPER, in the three biological replicates of both ds*U16*- and ds*GFP*-injected samples. Names of HdIV loci are indicated as well as the scaffold (Scaf-) they are located in. **(C)** Coverage values for proviral segment loci in the three biological replicates of the ds*U16* and ds*GFP* samples. For better visualization, only the scaffold (Scaf-) in which the proviral segments are located is indicated. The list of the proviral segment loci within each scaffold is available in Table 1. The y-axis was transformed by the log function for better data visualization. Statistical analyses are available at https://github.com/flegeai/EVE_amplification.

We extended our analysis by injecting ds*GFP* or ds*U16* into newly formed pupae, followed by isolation of DNA from calyx cells and hind legs, where no HdIV replication occurs. We then used specific primers and qPCR assays that measured DNA abundance of three wasp genes, selected HdIV replication genes inside and outside of IVSPERs, and selected HdIV genes in different proviral segments. As anticipated, no genes were amplified in hind legs from either control or treatment wasps (Fig 6). In ds*GFP*-injected control wasps, all HdIV genes were amplified in calyx cell samples (Fig 6). Among the wasp genes, only *XRCC1* exhibited significant amplification, consistent with its location within the IVSPER-1 amplified region (Fig 6). In contrast, when examining calyx cell DNA from wasps injected with ds*U16*, none of the HdIV genes nor *XRCC1* were amplified (Fig 6). Altogether, our results indicated U16 is required for amplification of all HdIV loci.

**Fig 6.**
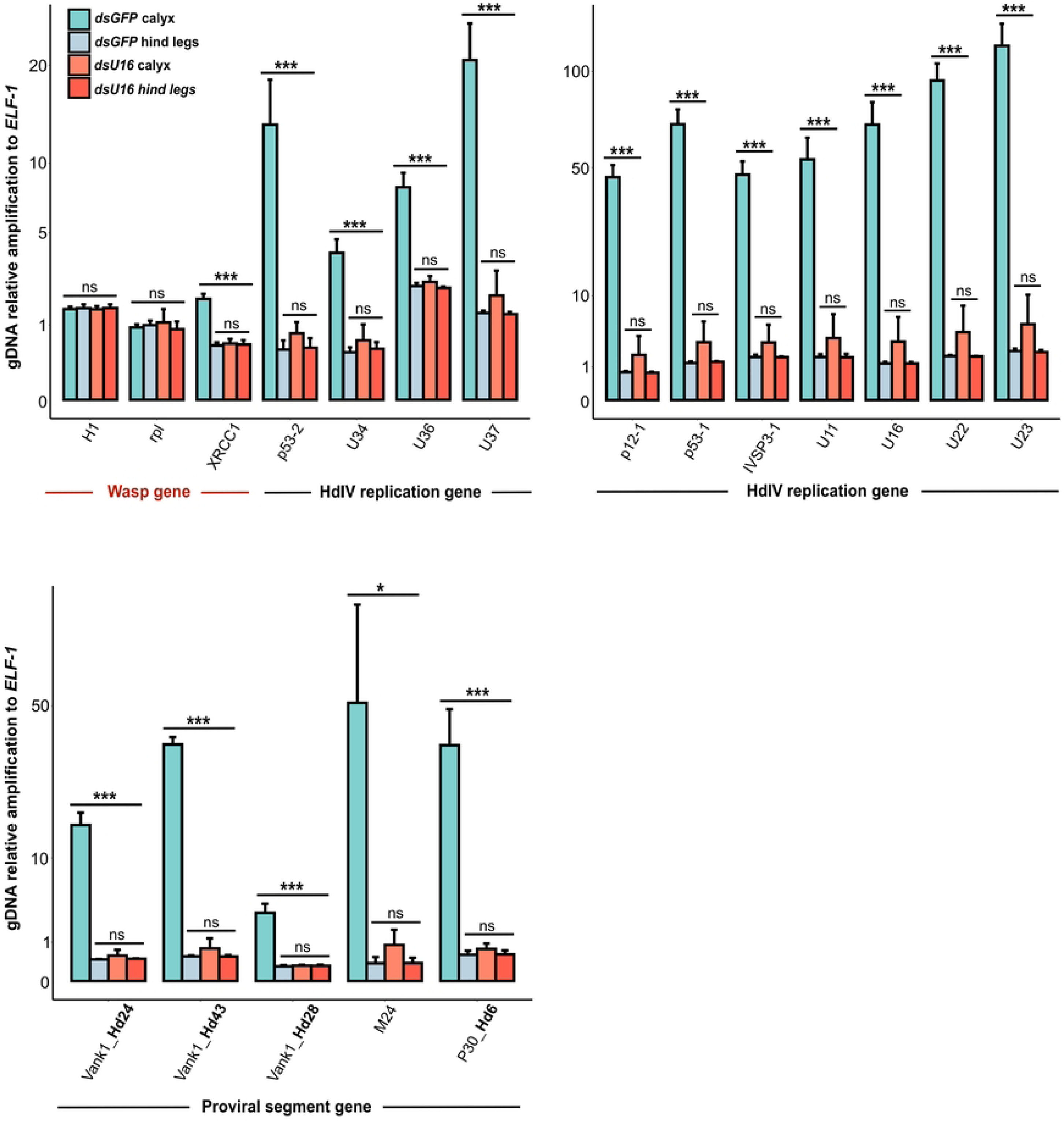
Impact of *U16* RNAi knockdown on amplification of select wasp and HdIV genes. Relative genomic amplification of selected HdIV genes in two-day-old females injected with ds*GFP* or ds*U16*. The wasp gene *XRCC1*, located within the amplified region of the IVSPER-1 locus, was incorporated into the analysis. Wasp histone (H1) and ribosomal protein (rpl) genes served as controls. Samples were obtained from calyx cells (where virion are produced) and hind legs (control). Statistical significance levels are denoted as follows: ns = non-significant, *p<0.05, **p<0.01, and ***p<0.001. The y-axis values were transformed using the square root function for better data visualization.

### Impact of DNA amplification on IV replication gene transcription levels and abundance of circularized HdIV molecules in calyx cells

We hypothesized that amplification of IV replication genes would increase transcript abundance which in turn would be affected by inhibiting HdIV DNA amplification. We thus compared transcript abundance of various genes in IVSPER-1, -2, and -3, in calyx RNA samples that were collected from wasps treated with ds*U16* or ds*GFP. U16* knockdown reduced expression of every HdIV replication gene we examined (Fig 7A). Finally, we investigated the impact of *U16* knockdown on the abundance of the circularized dsDNAs that are processed from amplified proviral segments. For this assay, we used PCR primers that specifically amplified the proviral form, circularized (episomal) form or both forms of Hd29 (Fig 7B). Results showed a significant reduction in both the proviral and circularized forms of Hd29 in calyx cell DNA from wasps injected with ds*U16* when compared to DNA from wasps injected with ds*GFP* (Fig. 7B). Our results thus indicated U16 is required for proviral segment amplification which is also required for production of circularized segments.

**Fig 7.**
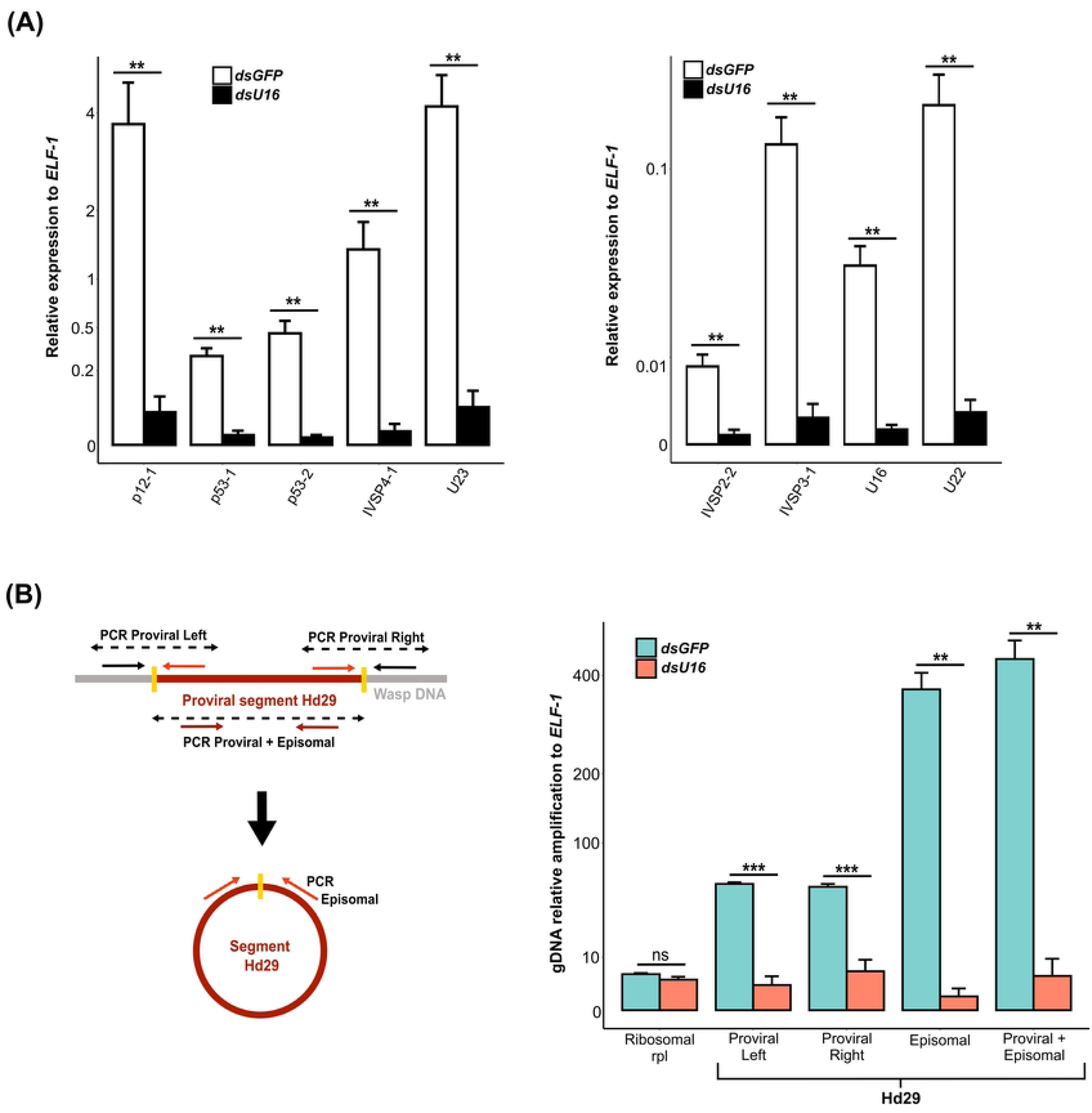
Impact of *U16* RNAi knockdown on HdIV replication gene expression and proviral segment amplification. **(A)** Relative expression of nine IVSPER genes in 2-day-old adult females injected with ds*GFP* (control) or ds*U16*. **(B)** Relative DNA amplification of the integrated linear (proviral) and circularized (episomal) forms of viral segment Hd29 in 2-day-old adult females injected with ds*GFP* (control) or ds*U16*. The left panel illustrates the position of primer pairs designed to selectively amplify the proviral form (Proviral Left and Right, indicated by red and black arrows), the circularized form (Episomal, red arrows), or both (Proviral + Episomal, brown arrows). The right panel presents the relative amplification of each form using DNA from ds*GFP*- and ds*U16*-injected females. In both **(A)** and **(B)**, significance levels are indicated as follows: ns = non-significant, *p<0.005, **p<0.01, and ***p<0.001. The y-axis values were transformed using the square root function for better data visualization.

## Discussion

During parasitism, wasps associated with IVs, BVs and other DEVs simultaneously inject virus-derived particles and eggs into their host. The role of DEV-derived particles in the success of wasp parasitism is well documented in the literature [22, 33, 34]. BVs, which evolved from a nudivirus, share a set of genes homologous to nudivirus and baculovirus core genes. Functional studies, guided in part by these similarities, have provided insights into several key processes underlying BV virion production. Identification of BV core genes that regulate the expression of other BV core genes encoding structural proteins [25], are involved in BV virion formation [25, 26, 35], or are required for processing proviral segments into circular DNA molecules packaged into capsids [25, 26] have been documented. In contrast, identifying the components of IV genomes and functions of IV genes regulating replication is more challenging because the hypothesized NCLDV ancestor is unknown. In turn, IV genome components with known or hypothesized functions in replication share little or no homology with known viruses. This study significantly advances understanding of IV replication by generating a chromosome level assembly for the *H. didymator* genome, presenting several lines of evidence showing that all HdIV loci are amplified in calyx cells when virions are being produced, and identifying *U16* as an essential gene for amplification of all HdIV loci and virion formation. This study also highlights the critical role of viral DNA amplification for IV virion production.

Earlier studies suggested IV proviral segment loci undergo amplification before viral segment processing [18, 19]. Another study indicated amplification of a few IVSPER genes and one proviral segment located in close vicinity of an IVSPER in one-day-old *H. didymator* adults [12]. However, the question persisted regarding whether all IV genome components were amplified in calyx cells and when amplification initiates during the time-course of virion production. To address these questions, we used our new chromosome-level genome assembly to map domains that undergo amplification in calyx cells during virion morphogenesis. Read mappings to genomic DNA extracted from *H. didymator* pupal stages 1 and 3 revealed that all HdIV genome components are simultaneously and locally amplified in calyx cells in stage 3 pupae. This analysis further identified five proviral segments and five IV replication genes located outside of IVSPERs that were previously unknown, resulting in a total of 67 HdIV proviral loci dispersed among the 12 *H. didymator* chromosomes. To elucidate the time-course of HdIV loci replication, the amplification of a subset of IV genome components was analyzed by qPCR. Our results show that HdIV loci amplification initiates between stage 1 and stage 2 pupae and reaches its maximum in stage 4 pupae. The temporal pattern observed in *H. didymator* is similar to BV-associated braconids. In the braconid wasp *Chelonus inanitus*, where the amplification kinetics of two proviral segments have been studied, local chromosomal amplification does not occur in the initial stages of pupal development [36]. Instead, it is preceded by an increase in DNA content through endoreduplication [37]. The question of whether calyx cell nuclei undergo polyploidization before local DNA amplification occurs in the case of *H. didymator* has yet to be investigated. Collectively, our results indicate DNA amplification of IV genome components constitutes one of the initial steps of virion morphogenesis.

Our data indicate all HdIV loci and genes located outside of IVSPERs are amplified with non-discrete boundaries that extend variable distances into flanking wasp DNA. In contrast to certain integrated viruses, such as polyomaviruses, which can be amplified in an “onion skin” type of replication with replication forks terminating at discrete boundaries [38], IVSPER amplification more closely resembles the local amplification observed in *Drosophila* follicle cells. In *Drosophila*, six loci corresponding to chorion genes or genes related to oogenesis are amplified in large regions of about 100 Kbp beyond the genes themselves, without discrete termination sites [39, 40]. Similar to IVSPERs, levels of DNA amplification in *Drosophila* follicle cells vary among different amplicons [40, 41]. In *Drosophila* follicle cells, amplification of these loci is associated with repeated firing of origins of replication (ORs) interspersed within each gene cluster. This results in overlapping bidirectional replication forks progressing outward on either side of the ORs [41]. These similarities between the pattern of DNA amplification of *Drosophila* genes and *H. didymator* proviral loci suggest that IVSPERs and IV replication genes may also be amplified through repeated firing of ORs present within the loci. However, additional approaches, such as nascent strand sequencing based on λ-exonuclease enrichment [42], will be necessary to identify ORs within IV genome components and validate this hypothesis.

Amplification of proviral segment loci is further characterized by a significant increase in read coverage at the Direct Repeat (DR) positions bordering the proviral segments, which serve as sites for homologous recombination and circularization of the segments. This suggests that a portion of the rapid increase in read coverage is due to reads mapping to amplification intermediates and circularized segments. The presence of circular forms in the sequenced genomic DNA samples is supported by our qPCR results for segment Hd29, which indicate the presence of amplicons specific to the circular form of Hd29 (Fig 7B). Accurately quantifying the proportion of reads mapping to the chromosomal form of HdIV segments, and estimating the actual extent of local DNA amplification presents a challenge. This is because paired-end reads that align within HdIV segment loci cannot discriminate between chromosomal HdIV DNA, potential replication intermediates, or circularized DNA. Nevertheless, considering the observed pattern of amplification in regions containing both IVSPERs and segments (Fig 3B), we propose that proviral segment loci may undergo amplification similar to IVSPERs or HdIV replication gene loci. The question persists regarding the subsequent processing of chromosomally amplified DNA and the mechanism behind the generation of a large number of circular molecules. The short-read data generated in this study have several limitations in characterizing whether amplification of proviral segment loci generates concatemeric intermediates and, if so, their orientation. Long-read data will be necessary to address these questions. Nonetheless, our results suggest HdIV proviral segment amplification involves both local chromosomal amplification and amplification of intermediates related to producing the circular dsDNAs that are packaged into capsids.

Our interest in *U16* stemmed from previous results indicating it is transcriptionally upregulated in calyx cells before the appearance of envelope and capsid components [16]. Sequence analysis during this study revealed a PriCT-2 domain in U16, known from primases in herpesviruses, whose function is unknown but may facilitate the association of the large primase domain (AEP) with DNA [31, 43]. Although other known primase domains were not identified in the U16 sequence, the presence of a PriCT-2 domain suggested this protein might play a role in the replication of HdIV genome components. Additionally, our RNAi experiments demonstrate that *U16* knockdown resulted in the complete absence of virion production in calyx cell nuclei and calyx fluid. These observations indicated an essential role for U16 in the early stages of viral replication, potentially involved in the amplification of HdIV genome components and/or the transcriptional regulation of IV replication genes. Subsequently, we analyzed the genome-wide impact of RNAi knockdown of *U16* on HdIV loci amplification, revealing that this gene is crucial for the amplification of all *H. didymator* IV genome components. In the case of IV replication genes, reduced amplification was accompanied by a simultaneous significant reduction in transcript abundance, likely resulting in insufficient amounts of HdIV structural proteins. However, amplification and transcription abundance levels did not fully correlate with each other. For instance, *U11* and *IVSP3-1* (both located on IVSPER-2) exhibit similar amplification patterns (Fig 1), but earlier findings showed that transcript abundances were not the same in calyx cells [15]. Thus, differences in gene expression observed among genes located within the same amplified regions (Fig 1) could also be affected by promoter strength or other factors. On the other hand, inhibition of proviral segment loci amplification had consequences for the abundance of the circularized dsDNA that are packaged into capsids, which were drastically reduced. Thus, our results identify U16 as an essential protein for virion morphogenesis. However, its precise role in viral replication remains to be understood. Questions to be addressed in the future include whether U16 acts at the initiation or elongation step of HdIV DNA replication, whether it interacts directly with DNA, or with proteins from the replisome complex, which itself could be composed of a mixture of HdIV and wasp proteins.

BVs share some features with IVs but also exhibit differences. Notably, in contrast to IVs, where most core genes with functions in virion morphogenesis reside in IVSPERs, many BV core replication genes are widely dispersed in the genomes of wasps [44, 45, 46] and are not amplified in calyx cells during virion morphogenesis [47]. However, the genomes of some BV-producing wasps do contain a ∼400 kb DNA domain in which several nudiviral core genes are located, known as the nudivirus-like cluster. This feature potentially identifies a site where the nudivirus ancestor of BVs integrated into the common ancestor of microgastroid braconids [9]. Notably, the nudivirus-like cluster is amplified with non-discrete boundaries [47], similar to what is reported for IV genome components in this study. The observed similarity in the amplification pattern between the BV nudivirus cluster and the proviral components of IVs could suggest they are amplified through a common mechanism, even though the molecules involved differ.

BV genomes also contain proviral segment loci with boundaries defined by flanking DRs and amplified in regions that include flanking regions outside of each DR. However, unlike IV proviral segments, the amplified flanking regions in BVs contain very precise nucleotide junctions that identify the boundaries of amplification [47, 48]. It is also known that some BV proviral segments are amplified as head-to-tail concatemers, consistent with a rolling circle amplification mechanism, while others are amplified as head-to-head and tail-to-tail concatemers, suggesting amplification by different mechanisms. However, all of these concatemers are similarly processed into circular DNAs by recombination at a precise site within DRs, which is a tetramer conserved in all BV segments [47, 48]. Nudiviral genes encoding tyrosine recombinases are further known to mediate this homologous recombination event [25, 26]. These types of molecules could also be present in IV genomes and need to be discovered. Currently, a detailed comparison between BV and IV proviral segment amplification is challenging and will require more information about the machinery involved in the processing of IV proviral segments into circular dsDNAs that are packaged into capsids.

Collectively, our results identify *U16* as a gene deriving from the IV ancestor that is required for HdIV DNA replication. This suggests that viral regulatory factors required for DNA amplification other than U16 have been preserved in parasitoid genomes. U16 may also interact with wasp cellular machinery in regulating DNA amplification, virion morphogenesis or both. Furthermore, this work emphasizes the value of studying original endogenized viruses, such as those found in parasitoids, to unveil new regulators of DNA processing.

## Materials and Methods

### Insects

*H. didymator* was reared as previously outlined by [49]. Female pupae obtained from cocoons were staged using pigmentation patterns: stage 1, corresponding to hyaline pupae (approximately 3-day-old pupae); stage 2, had a pigmented thorax (4-day-old); stage 3, had a pigmented thorax and abdomen (5-day-old); stage 4, were pharate adults just before emergence.

### Dovetail Omni-C Library Preparation and Sequencing

DNA from 10 male offspring (i.e., haploid genomes) from a single female *H. didymator* was sent on dry ice to Dovetail Genomics for Omni-C™ library construction. In the process of constructing the Dovetail Omni-C library, chromatin was fixed in place within the nucleus using formaldehyde and subsequently extracted. The fixed chromatin was digested with DNAse I followed by repair of chromatin ends and ligation to a biotinylated bridge adapter. Proximity ligation of adapter-containing ends ensued. Post-proximity ligation, crosslinks were reversed, and the DNA was purified. The purified DNA underwent treatment to eliminate biotin not internal to ligated fragments. Sequencing libraries were generated utilizing NEBNext Ultra enzymes and Illumina-compatible adapters. Fragments containing biotin were isolated using streptavidin beads before PCR enrichment of each library. The library was sequenced using the Illumina HiSeqX platform, which generated approximately 30x coverage. Subsequently, HiRise utilized reads with a mapping quality greater than 50 (MQ>50) for scaffolding purposes.

### Scaffolding the Assembly with HiRise

The *de novo* assembly from [15], and the Dovetail OmniC library reads served as input data for HiRise, a specialized software pipeline designed for leveraging proximity ligation data to scaffold genome assemblies, as outlined by [50]. The sequences from the Dovetail OmniC library were aligned to the initial draft assembly using the bwa tool (available at https://github.com/lh3/bwa). HiRise then analyzed the separations of Dovetail OmniC read pairs mapped within the draft scaffolds. This analysis generated a likelihood model for the genomic distance between read pairs. The model was subsequently employed to identify and rectify putative misjoins, score potential joins, and execute joins above a specified threshold. A contact map was generated from a BAM file by utilizing read pairs where both ends were aligned with a mapping quality of 60.

### Genomic DNA (gDNA) extraction for high throughput sequencing

Comparative analysis of two pupal stages. Genomic DNA (gDNA) was extracted from pooled calyx samples dissected from H. didymator female pupae at stage 1 (∼60 females) and stage 3 (∼50 females). Since the aim was to compare the two developmental pupal stages, a single replicate was done for each stage. Impact of U16 knockdown. Genomic DNA from calyces was collected from stage 3 female pupae that were injected with dsGFP and dsU16. This experiment involved three biological replicates, each corresponding to 30 to 50 calyx samples. Genomic DNA was extracted using the phenol-chloroform method. Briefly, calyx samples were incubated in proteinase K (Ambion, 0.5 μg/μl) and Sarkosyl detergent (Sigma, 20%), followed by treatment with RNAse (Promega, 0.3 μg/μl). Total genomic DNA was then extracted through phenol-chloroform extraction and ethanol precipitation. Following extraction, gDNA was quantified using a QBIT fluorometer (ThermoFisher) and subsequently sent for sequencing to Genewiz/Azenta company. Paired-end sequencing was carried out using Illumina technology and NovaSeq 2x150bp platform.

### NGS data analyses

Illumina reads were aligned to the updated version of the *H. didymator* genome using bwa mem [51], version 0.7.17, with default parameters. Subsequently, the aligned reads were converted to BAM files utilizing samtools view (version 1.15) [52].

*Prediction of the amplified regions.* Amplification peaks were identified using MACS2 [30] by comparing the pupal stage 3 alignment file as treatment and the pupal stage 1 alignment file as control. The specified parameters for this analysis were: --broad --nomodel -g 1.8e8 -q 0.01 --min-length 5000. Out of the 165 predicted peaks (i.e., amplified regions), only those with a fold change (FC) higher than 2 were retained for further analyses, resulting in a total of 59 peaks. These 59 peaks encompassed all known proviral loci, except for Hd40, which had a slightly lower value than the specified threshold (FC=1.9), and Hd45.1 and Hd2-like, located too close to the scaffold end and potentially missed. For the predicted peaks with FC>2 that did not correspond to known proviral loci, a manual curation was performed to determine whether these regions corresponded to HdIV loci. Proviral segments were identified by their flanking direct repeats (DRs) and gene contents, specifically the presence of genes belonging to IV segment conserved gene families. To identify putative core IV replication genes, genes present in the MACS2 peak were analyzed. Only those with no similarity to wasp proteins and that were transcribed in calyx cells (based on the available transcriptome from [16]) were retained.

*Read coverage per proviral region (HdIV locus or amplified region).* Raw read counts were determined for each proviral region using featureCounts [53] from the Subread package (version 2.0.1) with the parameters (-c -P -s 0 -O). Subsequently, coverage values were computed with a custom script available at https://github.com/flegeai/EVE_amplification. Coverage values for each region were calculated by dividing the number of fragments mapped to the region by the size of the region (expressed in kilobase pairs, kbp), and further normalized by the depth of the library (expressed in million reads). These coverages were computed for various types of genomic regions, including each locus (IVSPERs, IV replication genes outside IVSPERs, proviral segments), each MACS2-detected amplified region, and for each pupal stage (stage 1, St1 and stage 3, St3), as well as for each experiment (ds*GFP* and ds*U16*) and each replicate.

*Genome coverages per position on* H. didymator *scaffolds* (Counts per Million, CPM) and *Maximal value of amplification per proviral locus*. Genome coverages per position in 10 bp bins were acquired using the BamCoverage tool from the deeptools package [54] with the options: --normalizeUsing CPM and - bs 10. Subsequently, for each 10 bp bin, the pupal stage 3 (St3) versus stage 1 (St1) ratio was computed through an in-house script available at https://github.com/flegeai/EVE_amplification. This script utilized the pyBigWig python library from deeptools [54]. To determine the maximal counts per million (CPM) at each stage for every proviral locus, an in-house script importing the pyBigWig python library was employed. The maximum CPM value for the “stage 3 / stage 1” ratio was then calculated based on the 10 bp bin bigwig file, specifically for the position displaying the highest CPM value at stage 3 (summit).

*Comparison of read coverages between HdIV loci and the rest of the wasp genome.* One hundred sets of random regions, each mimicking the size distribution of HdIV loci, were generated using the shuffle tool from bedtools version 2.27 [55]. This was achieved by utilizing the bed file of HdIV loci (56 for proviral segments and 11 for IVSPERs) as parameters for the shuffle tool. Raw read counts for these randomly generated regions were computed in the same manner as for proviral regions, employing featureCounts [53] from the Subread package (version 2.0.1) with the parameters (-c -P -s 0 -O). Subsequently, coverage values per region were calculated using the same methodology as described earlier, with an in-house script available at https://github.com/flegeai/EVE_amplification.

*Search for motifs at the HdIV amplified regions boundaries.* The MEME suite [56] was employed for analyses using default parameters and a search for six motifs. A dataset comprising a total of 110 sequences, each spanning 1,000 nucleotides on both sides of the start and end positions of the 55 HdIV amplified regions predicted by the MACS2 algorithm, was utilized for this analysis. As a control, a parallel analysis was conducted using 110 sequences, each 2,000 nucleotides in length, randomly selected from locations within the *H. didymator* genome but outside the HdIV loci. This control dataset allowed for the comparison of motif patterns between the HdIV amplified regions and randomly chosen genomic regions.

### Genomic DNA extraction for gDNA amplification analyses by quantitative real-time PCR

To assess the level of DNA amplification, total genomic DNA (gDNA) was extracted using the DNeasy Blood & Tissue Kit (Qiagen) following the manufacturer’s protocol. Ovaries (ovarioles removed) and hind legs, representing the negative control, were dissected from ten pupae at four different stages. Three replicates were generated for each pupal stage. Quantification of target gene amplification was conducted through quantitative PCR, utilizing LightCycler® 480 SYBR Green I Master Mix (Roche) in 384-well plates (Roche). The total reaction volume per well was 3 µl, comprising 1.75 µl of the reaction mix (1.49 µl SYBR Green I Master Mix, 0.1 µl nuclease-free water, and 0.16 µl diluted primer), and 1.25 µl of each gDNA sample diluted to achieve a concentration of 1.2 ng/µl. Primers used are listed in S4 Table. The gDNA levels corresponding to the viral genes and the housekeeping wasp gene (elongation factor (ELF-1)) were determined using the LightCycler 480 System (Roche). The cycling conditions involved heating at 95°C for 10 min, followed by 45 cycles of 95°C for 10 s, 58°C for 10 s, and 72°C for 10 s. Each sample was evaluated in triplicate. The obtained DNA levels were normalized with respect to the wasp gene ELF-1. Raw data are provided in S3 Dataset.

### Total RNA extraction

Total RNA was extracted from ovaries (ovarioles removed) dissected from pupae at different stages using the Qiagen RNeasy extraction kit in accordance with the manufacturer’s protocol. To control for gene silencing, total RNAs were also extracted from individual adult wasp abdomens (2 to 4 days old). For this, Trizol reagent (Ambion) was initially used followed by extraction using the NucleoSpin® RNA kit (Macherey-Nagel). Isolated RNA was then subjected to DNase treatment using the TURBO DNA-free Kit (Life Technologies) to assure removal of any residual genomic DNA from the RNA samples.

### Protein sequence analyses

Conserved domains of U16 were identified using the CD-search tool available through NCBI’s conserved domain database resource [57, 58]. Subcellular localization predictions were made using the DeepLoc - 2.0 tool, a deep learning-based approach for predicting the subcellular localization of eukaryotic proteins [59]. For multiple sequence alignment, CLUSTAL Omega (version 1.2.4) was employed [60]. Structure predictions for U16 were carried out using the MPI Bioinformatics Toolkit [61].

### RNA interference (RNAi)

Gene-specific double-stranded RNA (dsRNA) used for RNAi experiments was prepared using the T7 RiboMAX™ Express RNAi System (Promega). Initially, a 350-450 bp fragment corresponding to the *U16* sequence was cloned into the double T7 vector L4440 (a gift from Andrew Fire, Addgene plasmid # 1654). Subsequently, an *in vitro* transcription template DNA was PCR amplified with a T7 primer, and this template was used to synthesize sense and antisense RNA strands with T7 RNA polymerase at 37°C for 5 hours. The primers used for dsRNA production are listed in S4 Table. After annealing and DNase treatment using the TURBO DNA-free Kit (Life Technologies), the purified dsRNAs were resuspended in nuclease-free water, quantified using a NanoDrop ND-1000 Spectrophotometer (Thermo Scientific), and examined by agarose gel electrophoresis to ensure their integrity. Injections were performed in less than one-day-old female pupae using a microinjector (Fentojet® Express, Eppendorf®) and a micromanipulator (Narishige®). Approximately 0.3-0.6 μl of 500 ng/μl dsRNA was injected into each individual. Control wasps were injected with a non-specific dsRNA homologous to the green fluorescent protein (GFP) gene. Treated pupae were kept in an incubator until adult emergence, which occurred approximately 5 days after injection.

### Transmission electron microscopy

Ovaries were dissected from adult wasps between 2 and 3 days after emergence, following the procedures outlined in [17]. To ensure consistency of the observed phenotype, at least three females (taken at different microinjection dates) were observed for each tested dsRNA. For transmission electron microscopy (TEM) observations, calyces were fixed in a solution of 2% glutaraldehyde in PBS for 2 hours and then post-fixed in 2% osmium tetroxide in the same buffer for 1 hour. Tissues were subsequently bulk-stained for 2 hours in a 5% aqueous uranyl acetate solution, dehydrated in ethanol, and embedded in EM812 resin (EMS). Ultrathin sections were double-stained with Uranyless (DeltaMicroscopy) and lead citrate before examination under a Jeol 1200 EXII electron microscope at 100 kV (MEA Platform, University of Montpellier). Images were captured with an EMSIS Olympus Quemesa 11 Megapixels camera and analyzed using ImageJ software [62].

### Reverse-transcriptase quantitative real-time PCR (RT-qPCR)

For RT-qPCR assays, 400 ng of total RNA was reverse-transcribed using the SuperScript III Reverse Transcriptase kit (Life Technologies) and oligo(dT)15 primer (Promega). The mRNA transcript levels of selected IVSPER genes were measured by quantitative reverse transcription-PCR (qRT-PCR) using a LightCycler® 480 System (Roche) and SYBR Green I Master Mix (Roche). Expression levels were normalized relative to a housekeeping wasp gene (elongation factor 1 ELF-1). Each sample was evaluated in triplicate, and the total reaction volume per well was 3 µl, including 0.5 µM of each primer and cDNA corresponding to 0.88 ng of total RNA. The amplification program consisted of an initial step at 95°C for 10 min, followed by 45 cycles of 95°C for 10 s, 58°C for 10 s, and 72°C for 10 s. The primers used for this analysis are listed in S4 Table.

### qPCR data analysis

Data were acquired using Light-Cycler® 480 software. PCR amplification efficiency (E) for each primer pair was determined by linear regression of a dilution series (5x) of the cDNA pool. Relative expression, using the housekeeping gene ELF-1 as a reference, was calculated through advanced relative quantification (Efficiency method) software provided by Light-Cycler® 480 software. For statistical analyses, Levene’s and Shapiro-Wilk tests were employed to verify homogeneity of variance and normal distribution of data among the tested groups. Differences in gene relative expression between developmental stages and between ds*GFP* and ds*U16*-injected females were assessed using a two-tailed unpaired t-test for group comparison. In cases where homogeneity of variance was not assumed, the Welch-test was used to compare gene relative expression between groups. A p-value < 0.05 was considered significant. All statistical analyses were conducted using R [63]. Detailed statistical analyses of qPCR results are provided in S3 Dataset.

### Data availability

The datasets supporting the conclusions in this article are accessible at the NCBI Sequence Read Archive (SRA) under the Bioproject accession number PRJNA589497. Additionally, the new version of the *H. didymator* genome, annotation, alignments of reads, and coverage information can be found at BIPAA (https://bipaa.genouest.org/sp/hyposoter_didymator/). Raw data and statistical analyses for all the qPCR analyses are provided in S3 Dataset. Furthermore, sequencing raw data, read coverage analyses, statistical analyses, and in-house scripts are available at https://github.com/flegeai/EVE_amplification.

## Acknowledgments

The insects used in the experiments were provided by Raphaël BOUSQUET and Gaétan CLABOTS from the DGIMI insect rearing facility. All RNAi experiments were conducted in the insect quarantine platform (PIQ) of DGIMI lab, which is a member of the Montpellier Vectopole Sud network (https://www.vectopole-sud.fr/). Microscopy observations were facilitated by the Montpellier MEA platform (https://mea.edu.umontpellier.fr/). All qPCR analyses were performed with the assistance of the Montpellier Genomix qPHD platform (http://www.pbs.univ-montp2.fr/).

## Supporting information captions

**S1 Dataset**. *Hyposoter didymator* Hi-C genome assembly. The dataset includes: **A.** Figure depicting the Hi-C scaffold contact map; **B.** Table presenting the Hi-C scaffolds containing HdIV loci; **C.** Figure displaying the pairwise comparisons of HdIV segments located in close proximity within the *H. didymator* scaffolds.

**S2 Dataset**. Sequence analysis and alignment of the U16 gene from *H. didymator* to four other wasp species that harbor IVs. The dataset includes: **A.** Multiple sequence alignment of U16 proteins from different parasitoid species. **B.** Detail of the predicted secondary structure of the PricT-2 domain in the*H. didymator* U16 protein. **C.** Subcellular localization of U16 predicted by DeepLoc 2.0.

**S3 Dataset.** Raw data and statistical analyses of qPCR analyses. The dataset includes raw data and statistical analyses for: **A.** Genomic DNA amplification of IVSPER genes at four different *H. didymator* pupal stages; **B.** Genomic DNA amplification of IVSPER and HdIV segment genes in ds*GFP* and ds*U16*-injected wasps; **C.** RNA quantification of IVSPER genes in ds*GFP* and ds*U16*-injected wasps; **D.** DNA amplification of Hd29 segment in ds*GFP* and ds*U16*-injected wasps.

**S1 Table.** Read coverage of HdIV loci on each scaffold of the *H. didymator* genome.

**S2 Table.** List of the peaks predicted in *H. didymator* genome scaffolds using MACS2 algorithm.

**S3 Table.** Read coverage of HdIV amplified regions in calyx cell DNA from ds*GFP*- and ds*U16*-injected female pupae.

**S4 Table**. List of primers used in the present work.

**S1 Fig**. DNA amplification patterns of HdIV loci in calyx cells of *H. didymator*.

**S2 Fig**. HdIV amplified regions in Scaffold-11.

**S3 Fig**. MEME analysis of boundaries of the predicted MACS2 HdIV amplified regions.

## Author contribution

A. LORENZI: Data curation, Formal analysis, Investigation, Methodology, Validation, Visualization, Writing – Original Draft Preparation, Writing – Review & Editing

F. LEGEAI: Data curation, Formal analysis, Investigation, Methodology, Validation, Visualization, Writing – Original Draft Preparation, Writing – Review & Editing

V. JOUAN, P.-A. GIRARD, M. EYCHENNE, M. RAVALLEC, Investigation, Methodology, Validation

A. BRETAUDEAU, S. ROBIN, Data Curation

J. ROCHEFORT, M. VILLEGAS, Investigation

M. R. STRAND, G. R. BURKE, Writing – Review & Editing

R. REBOLLO, Funding Acquisition, Validation, Writing – Original Draft Preparation, Writing – Review & Editing

N. NÈGRE, Conceptualization, Data curation, Funding acquisition, Investigation, Methodology, Resources, Supervision, Validation, Writing – original draft, Writing – review & editing

A.-N. VOLKOFF, Conceptualization, Data curation, Formal analysis, Funding acquisition, Investigation, Methodology, Project administration, Resources, Supervision, Validation, Visualization, Writing – original draft, Writing – review & editing

## Fundings

This work has been financially supported by the INRAE SPE department (EPIHYPO project) and the French National Research Agency (ENDOVIRE project, #ANR-22-CE20-0005-01). The Dovetail sequencing of the *H. didymator* genome has received funding from the European Union’s Horizon 2020 research and innovation program under the Marie Skłodowska-Curie grant agreement no. 764840 for the ITN IGNITE project, with Denis TAGU from IGEPP as a partner.

## Notes

### Competing Interest Statement

The authors have declared no competing interest.

